# Quantitative Mining of Compositional Heterogeneity in Cryo-EM Datasets of Ribosome Assembly Intermediates

**DOI:** 10.1101/2021.06.23.449614

**Authors:** Jessica N. Rabuck-Gibbons, Dmitry Lyumkis, James R. Williamson

## Abstract

Macromolecular complexes are dynamic entities whose function is often intertwined with their many structural configurations. Single particle cryo-electron microscopy (cryo-EM) offers a unique opportunity to characterize macromolecular structural heterogeneity by virtue of its ability to place distinct populations into different groups through computational classification. However, current workflows are limited, and there is a dearth of tools for surveying the heterogeneity landscape, quantitatively analyzing heterogeneous particle populations after classification, deciding how many unique classes are represented by the data, and accurately cross-comparing reconstructions. Here, we develop a workflow that contains discovery and analysis modules to quantitatively mine cryo-EM data for a set of structures with maximal diversity. This workflow was applied to a dataset of *E. coli* 50S ribosome assembly intermediates, which is characterized by significant structural heterogeneity. We identified new branch points in the assembly process and characterized the interactions of an assembly factor with immature intermediates. While the tools described here were developed for ribosome assembly, they should be broadly applicable to the analysis of other heterogeneous cryo-EM datasets.

## Introduction

Cryo-electron microscopy (cryo-EM) is a rapidly evolving, powerful technology for solving the structures of a wide variety of biological assemblies. The “resolution revolution” in cryo-EM (Kühlbrandt, 2014), caused in part by advances in direct electron detectors and improved data acquisition and analysis workflows, has led to high-resolution structural insights into a wide variety of biological processes performed by macromolecular assemblies (Fernandez-Leiro and Scheres, 2016). There have been steady, but consistent improvements to achievable resolution, and the collective tools are now enabling structure determination at true atomic resolution (Bartesaghi *et al.*, 2015; Tan *et al.*, 2018; Nakane *et al.*, 2020; Yip *et al.*, 2020; Zhang *et al.*, 2020). There have also been numerous advances in workflows for analyzing structurally heterogeneous particle populations, and data processing software now routinely include strategies for handling distributions of structures that arise from compositional or conformational changes in the macromolecular species of interest (Elmlund and Elmlund, 2012; Gao *et al.*, 2004; Klaholz, 2015; Liao, Hashem and Frank, 2015; Nakane *et al.*, 2018; Scheres, 2016; Spahn and Penczek, 2009; Wang *et al.*, 2013; White *et al.*, 2017; Zhong *et al.*, 2021; Grant, Rohou and Grigorieff, 2018; Lyumkis *et al.*, 2013; Punjani and Fleet, 2021b; Punjani and Fleet, 2021a). However, most current cryo-EM workflows still focus on achieving the maximum possible resolution, which requires selecting and averaging potentially heterogeneous subsets of the data in the interest of increasing the particle count for the homogeneous regions of a map. This strategy comes at the expense of either eliminating particle populations that do not conform to the predominant species or neglecting dynamic and labile regions of reconstructed maps, which are often of biological interest.

Another challenge in cryo-EM heterogeneity analysis is that there is no way to define the number of distinct structures in a given dataset *a priori*. It is up to the researcher to employ a classification strategy and to heuristically determine the number of distinct classes. Furthermore, there is no set procedure to determine the threshold for examining map features and differences between maps. Thresholds are often set in a subjective manner in order to best display the features of interest in the maps, although an approach was recently described where a voxel-based false discovery rate could be determined to establish a noise threshold for contouring (Beckers, Jakobi and Sachse, 2019). Thus, determining the final number of classes in a dataset and quantitatively comparing a set of maps in order to tell a concise biological story with statistical significance remains a challenge.

The process of ribosome assembly provides a useful case study for mining and quantitatively assessing structural heterogeneity in cryo-EM data. The bacterial 70S ribosome is a complex macromolecular machine composed of three ribosomal RNAs (rRNAs) and ~50 ribosomal proteins (r-proteins) that form a large 50S subunit and a small 30S subunit. Ribosome assembly occurs within several minutes *in vivo*, and the process includes transcription and translation of the rRNAs and r-proteins, folding of the rRNA and r-proteins, and docking of the r-proteins on the rRNA scaffold. rRNA folding events and proper r-protein binding are facilitated by ~100 ribosome assembly factors. Given the efficiency and speed of the assembly process, structural intermediates are difficult to isolate and purify. However, perturbations in ribosome assembly lead to the accumulation of numerous structural intermediates, which collectively inform molecular mechanisms of ribosome assembly (Shajani, Sykes and Williamson, 2011; Stokes *et al.*, 2014; Sashital *et al.*, 2014; Sykes *et al.*, 2010; Jomaa *et al.*, 2014; Li *et al.*, 2013; Ni *et al.*, 2016; Davis *et al.*, 2016; Rabuck-Gibbons *et al.*, 2020). The major parts of the ribosome that are often present or missing in assembly intermediates are the central protuberance (CP), the L7/12 and L1 stalks, and the base (Figure 1A).

**Figure 1.**
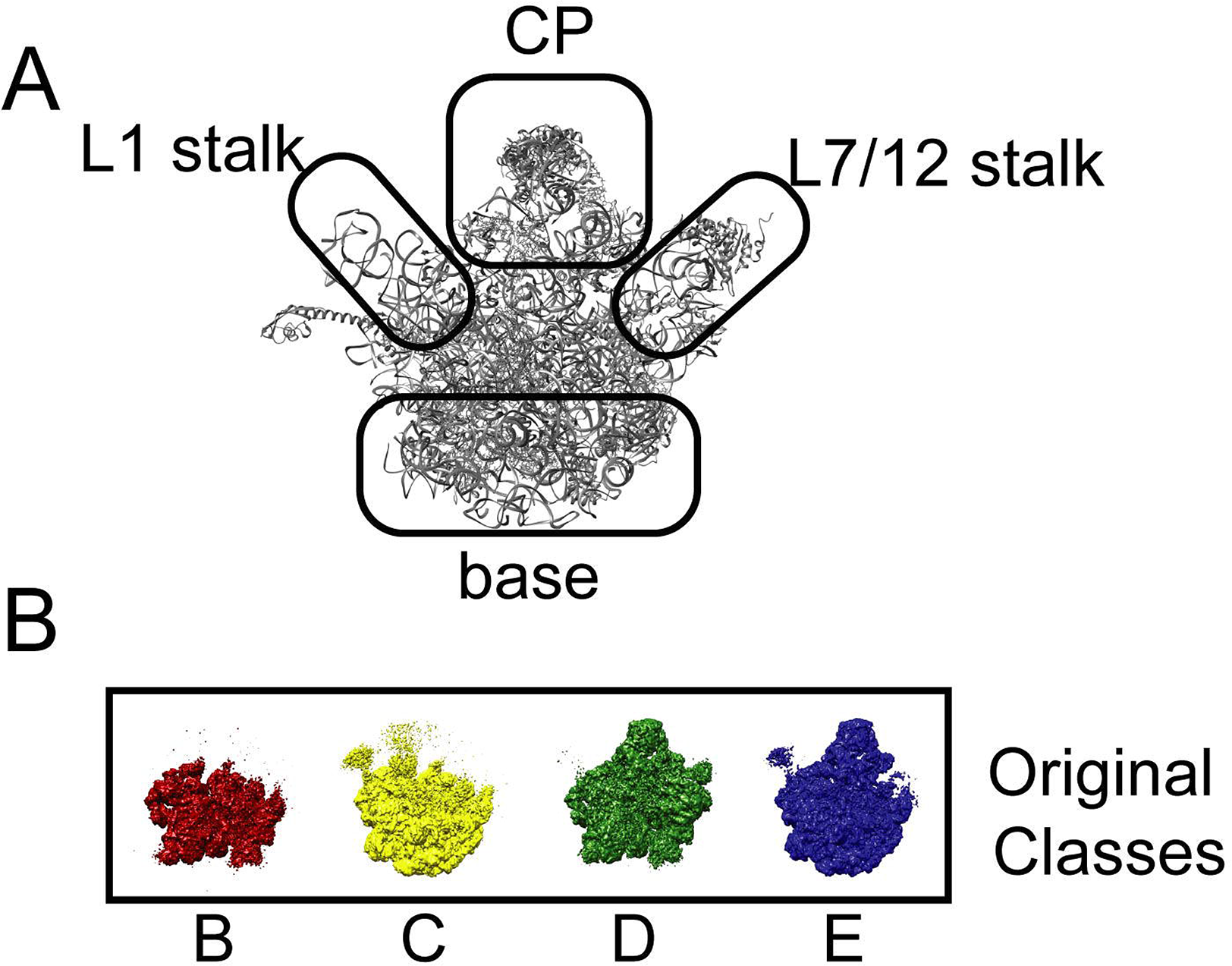
Description of the bacterial large ribosomal subunit and prior assembly intermediates identified by cryo-EM. (A) PDB ID 4YBB labeled with prominent features identifiable on the large ribosomal subunit, including the central protuberance (CP), base, L1 stalk, and L7/12 stalks. These terms are used throughout the paper. (B) Primary classes identified within the original bL17-lim dataset (Davis *et al.*, 2016). From left to right: B class (red), C class (yellow), D class (green), and the E class (blue).

We previously developed a genetic approach by which the amount of a given r-protein, in our case bL17, could be titrated by the addition of the small molecule homoserine lactone (HSL) (Davis *et al.*, 2016). Limiting the amount of bL17 induced a roadblock in ribosome assembly, causing intermediates to accumulate. In the first work using the bL17-lim strain (Davis *et al.*, 2016), we identified thirteen distinct structures that fell into four main structural classes (Figure 1B). Here, we will continue to use the nomenclature for the main classes used by Davis and Tan, et al. These categories, ordered least to most mature, are the B class which is missing the base, CP, and both stalks, the C class in which the base is formed, but the central protuberance (CP) is either misdocked or altogether missing, the D class in which the base of the 50S ribosome is missing, and the E class, which contains both the base and the CP, but has variability in the presence or absence of the stalks. Some of the “missing” regions (primarily rRNA, but they may also include r-proteins) described above are not present in the reconstructed maps but are in fact present in the sample and within individual particle images, meaning that they contribute to “biological noise”. This becomes relevant for some of the decisions that need to be made in the data analysis workflow, as will be discussed below. In previous work, the four main initial classes belonging to the 50S assembly intermediates (B, C, D, E) were each further subdivided by an additional round of subclassification, resulting in thirteen distinct structures. While several different subclassification schemes were attempted at that time using heuristics to determine the number of subclasses, no attempt was made to establish quantitative criteria by which the subclassification or coverage of relevant classes would be complete. While classes were identified belonging to the 30S and 70S (F class and A class in Davis *et al.*, 2016), they are not explicitly described in our previous work or in the work described here.

As our goal is to define broad trends in ribosome assembly through various perturbations, it is important to quantitatively assess differences between intermediates that accumulate under various specific conditions and to organize them into a ribosome assembly landscape (Davis *et al.*, 2016; Bernstein *et al.*, 2004; Harnpicharnchai *et al.*, 2001; Jomaa *et al.*, 2011; Loerke, Giesebrecht and Spahn, 2010; Nikolay *et al.*, 2018; Razi, Guarné and Ortega, 2017; Uicker, Schaefer and Britton, 2006). To this end, we developed a data processing framework to analyze cryo-EM datasets methodically and quantitatively in order to assess the number of distinct structures, the significant differences among them, and to place these structures into a biological context. When we apply our complete workflow to a dataset of ribosome assembly intermediates from bL17-lim, we discover a total of forty-one different structures that are identifiable based on a defined set of cutoff parameters. These structures include several novel intermediates, such as the most immature assembly intermediate observed to date, and an independent pathway contingent on the binding of a ribosome assembly factor, as well as late-stage assembly intermediates. Together, these are organized into a revised assembly landscape for the 50S ribosomal subunit under bL17-lim conditions.

## Results and Discussion

### An overview of the heterogeneity processing workflow

There are two main phases in the framework for systematic analysis of heterogeneous ensembles of macromolecular conformations (Figure 2). The first phase is a discovery phase, which begins with iterative rounds of hierarchical classification and sub-classification using a defined set of thresholding parameters. The goal of this first phase to uncover the broad spectrum of distinct classes in a cryo-EM dataset, starting with traditional pre-processing and data cleaning steps (e.g. motion correction, particle picking, CTF estimation, and initial 2D and 3D classification). The initial data cleaning steps defined here are intended to be very lenient, such that the only particles removed from the dataset are clear artifacts or molecular species that are not of interest. For example, in the case of 50S ribosome assembly intermediate analysis, we remove particles that are obvious 30S or 70S ribosomes and proteasomes from the stack, but we do not remove any classes that could possibly be 50S assembly intermediates. After the cleaning steps, an iterative subclassification strategy is used to parse out molecular heterogeneity. After an initial round of classification, each class (class X) is subjected to a n=2 subclassification, resulting in two potential subclasses, X1 and X2. Both subclasses are processed and binarized, and then difference maps X1-X2 and X2-X1 are calculated, to determine if there is more heterogeneity that can be mined from each class X. If the difference volumes don’t reach a chosen molecular weight or resolution threshold, then the subclassification is rejected, and further subclassification is terminated. If neither of these two criteria are reached, the binary subclassification process is iteratively repeated until one of the convergence criteria are met.

**Figure 2.**
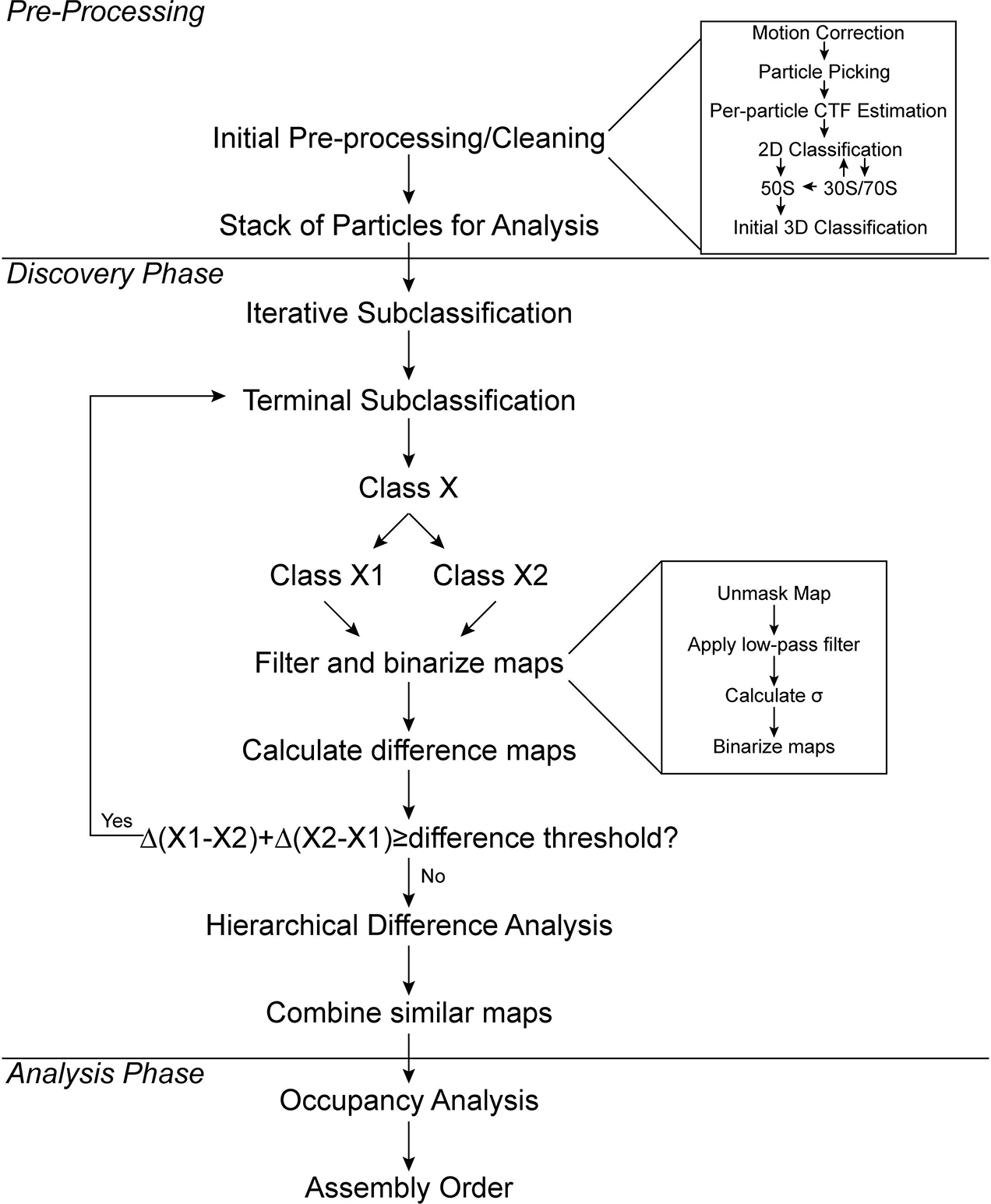
Workflow for cryo-EM heterogeneity analysis.

The second phase is an analysis phase, which is intended to quantitatively define and distinguish structural features between maps, and further, to establish the number of structural states using a given set of quantitative cutoffs. During this hierarchical difference analysis, the full matrix of difference maps is calculated, and the molecular weights of the difference maps are used as a metric to cluster the classes, which can be visualized as a particle dendrogram. A line can be drawn through the dendrogram at a chosen molecular weight threshold, which identifies similar maps that can be combined. Next, to qualitatively differentiate between classes, the resulting set of maps are compared to a catalog of coarse-grained structural features that are calculated from a reference structure, in this case the bacterial 50S ribosome. It is convenient to use features such as rRNA helices and r-proteins, that may be present or absent in various classes. The presence of these coarse-grained reference features is quantitatively analyzed using hierarchical clustering to organize and visualize the patterns of variation among the final set of particle classes. For our dataset of bacterial ribosome assembly intermediates, these features are used to place the observed classes into a putative assembly pathway, based on a principle of parsimonious folding and unfolding.

### A divisive resolution-limited subclassification approach facilitates identifying novel species

A major challenge in the analysis of heterogeneous datasets is the accurate identification of a broad diversity of structural states. To address this, we developed a classification strategy to mine an experimental cryo-EM dataset for distinct particle populations. Classification and refinement of particle classes can be undertaken using a variety of software packages, and we have adopted the latest version of FrealignX, whose code base is also implemented within *cis*TEM (Grant, Rohou and Grigorieff, 2018; Lyumkis *et al.*, 2013). We note that most processing packages that are capable of classifying single-particle cryo-EM data can be employed for this purpose(Scheres, 2016; Nakane *et al.*, 2018; Zhong *et al.*, 2021; Punjani and Fleet, 2021b; Punjani and Fleet, 2021a).

In typical cryo-EM workflows, 3D classification is performed several times, with different choices for the total number of classes (*n*). If *n* is too small, the resulting classes may have averaged properties leading to loss of structural diversity but potentially higher resolution in the homogeneous regions. If *n* is too large, the data is subdivided into nearly identical classes, but each class is characterized by lower resolution, because the particle count contributing to the class decreases. For the characterization of intrinsically heterogeneous datasets such as those encountered during ribosome assembly, the goal of 3D classification is to capture the full range of structural diversity, as opposed to a select few well-resolved classes. Therefore, we developed an iterative subclassification strategy to systematically mine the data and identify distinct structural intermediates, including species that are rare and underpopulated.

With the knowledge that our test dataset harbored at least thirteen intermediates (Davis *et al.*, 2016), we started with *n=*10 in order to evaluate parameters for subclassification. The ten initial classes are shown in Figure 3A. While we expected that we would find the previous B, C, D and E classes in the dataset, the B-class was not present, and rather, multiple classes that are subtle variations of the E-class were present. This exemplifies one of the pitfalls of classification that we term “hiding”, where subclasses can be mixed, only to emerge at subsequent stages of subclassification. A survey of various classification parameters within FrealignX revealed that lowering the *res_high_class* parameter, which is the resolution of the data to be used for classification, ameliorated class hiding and had a strong effect on the classes that emerged. This parameter is typically set to just below the estimated resolution limit of the data. However, by setting *res_high_class* to 20Å, the gross class heterogeneity increased, and the expected B-class emerged (Figure 3B). The resolution threshold for classification is frequently defaulted and determined automatically during classification, but it may also be explicitly set by the user or limited to the resolution of the first Thon ring (Scheres, 2012; Scheres, 2016; Scheres *et al.*, 2008). With the well-defined ribosome assembly case study, we show that a lower resolution threshold during classification helps to identify particle subsets that are substantially distinct from the predominant species.

**Figure 3.**
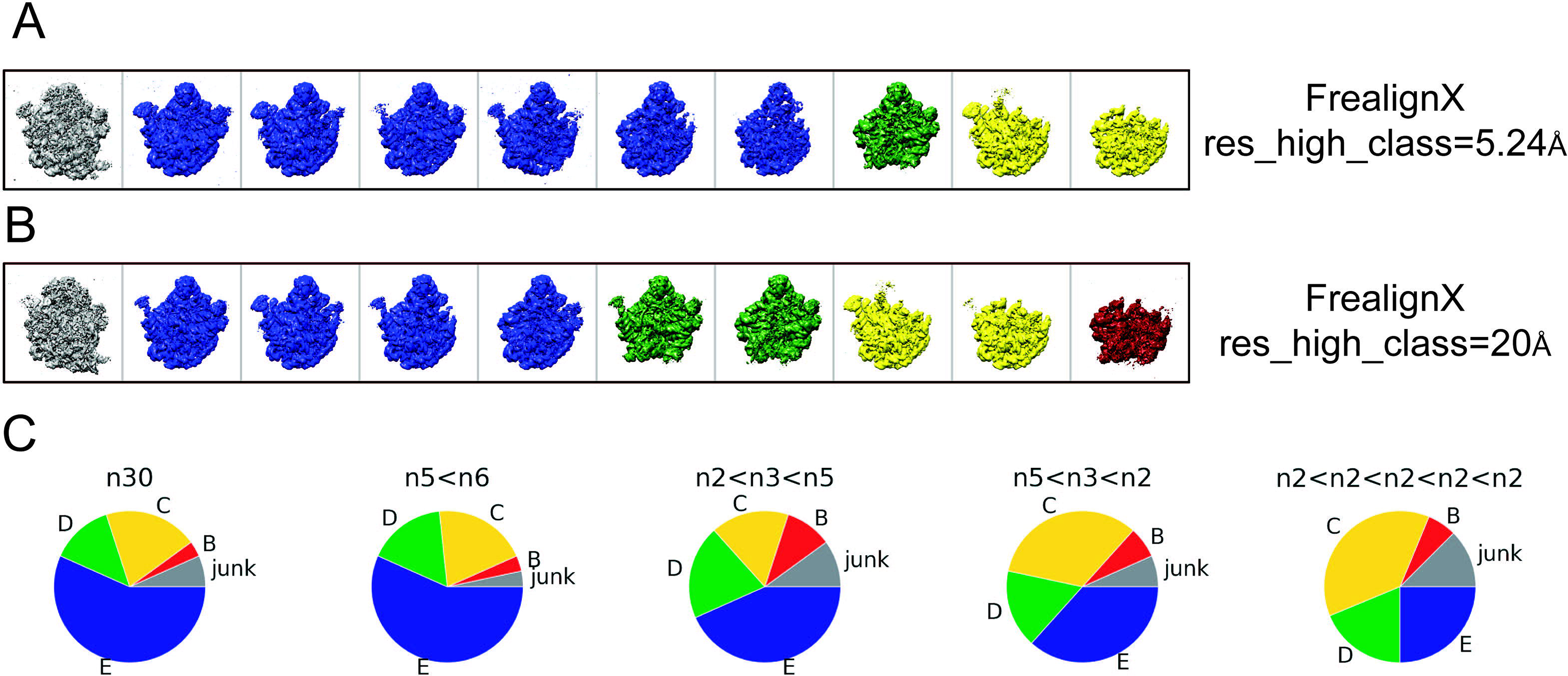
A divisive resolution-limited subclassification approach facilitates identifying rare structural variants. (A) FrealignX classification with *res_high_class* parameter set to Nyquist (5.24Å). (B) FrealignX classification with the *res_high_class* parameter set to 20Å. Using a lower resolution cutoff leads to the identification of a broader range of classes. (C) Results of the five different classification schemes. The colors (A,B) correspond to the classes found in (C).

We also examined different iterative subclassification strategies, with various numbers of classes used for each stage of subclassification. In order to test the success of these strategies, we selected a final *n* of ~30, which was chosen because it provided a convenient number to evaluate a variety of subclassification schemes, and because it was close to twice the final number of classes found in the original bL17-lim dataset (Davis *et al.*, 2016). The five classification schemes (Figure 3C) tested were: (1) a simple 1-round classification with *n*=30, (2) a 2-round hierarchical classification of 6 initial classes, each subdivided into 5 (n_1_=6,n_2_=5; total n=30,), (3) a 3-round hierarchical subclassification of 2 initial classes each subdivided into 3, with a second subdivision into 5 (n_1_=2,n_2_=3,n_3_=5; total n=30), (4) a 3-round hierarchical classification of 5 initial classes subdivided into 3, then subdivided into 2 n_1_=5,n_2_=3,n_3_=2; total n=30,), and (5) a 5-round hierarchical binary subclassification strategy, where 2 initial classes were subdivided into 2 until n=32 was reached (n_1_=2,n_2_=2,n_3_=2,n_4_=2,n_5_=2; total n=32,).

With the sole exception of the simple single-round classification with *n*=30, all of these divisive schemes yielded new classes not previously identified (Supplemental Figure 1, indicated by *). Furthermore, the iterative divisive approaches produced the greatest range of structural diversity and avoided grouping together dissimilar classes. This observation is perhaps not unexpected, as it is well known that a divisive classification approach avoids local minima within the search space and is more robust than attempting to produce a final number of classes directly (Gray, 1984; Sorzano *et al.*, 2010). Qualitatively, a first round of classification where n_1_ is on the order of the number of major classes works well, followed by smaller subdivisions. As an example, a three round subclassification scheme with [n_1_ = 5, n_2_ = 3, n_3_ = 2], for a total of 30 final classes, identified the greatest number of new structures, as shown in Supplemental Figure 1. For this reason, we proceeded with the n_1_ = 5, n_2_ = 3, n_3_ = 2 approach for our work, although we note that the optimal classification scheme will likely vary with the distinct heterogeneity spectrum for each unique dataset. Given that the observed classes are relatively independent of the details of the subclassification, we turned our attention to the criteria for termination of subclassification.

### Defining an endpoint for subclassification

The determination of when subclassification is complete is a key question in cryo-EM analysis. Many times, classification is considered finished if a specific region of interest can be resolved to a satisfactory resolution, depending on what question(s) the user wishes to address. However, this subjective approach may be insufficient for the purpose of uncovering hidden features and discovering new structural states, especially if there are multiple datasets to be compared. To guide the analysis of our bL17-lim dataset, and to establish a protocol that can be used to analyze other data with statistical significance, our goal was to establish metrics by which we could confidently terminate the subclassification. We adopted a simple metric to determine the endpoint of subclassification. For any given class at any stage of subclassification, a test subclassification is performed with n=2. If the two resulting subclasses differ by less than a chosen noise threshold, or by less than a chosen molecular weight threshold, then subclassification is complete, and the subdivision is rejected. Conversely, if the thresholds are exceeded, the subclassification is retained, and the two resulting classes are iteratively subjected to additional subclassification until the termination thresholds are met (Figure 2).

There are at least two types of noise that need to be considered in the difference analysis that are used to conclude subclassification. First, there is the intrinsic noise floor in the map that arises from averaging noisy image data during the reconstruction process. Second, there is biological noise, which can be broadly attributed to conformational and compositional heterogeneity, resulting in density above the intrinsic noise floor that cannot be interpreted in terms of a structure or slight shifts of well-defined elements that may or may not be significant (Supplemental Figure 2). For example, in the case of ribosome assembly, there are portions of rRNA that are present in the sample, but do not resolve to a reasonable structure (Davis *et al.*, 2016). To characterize a diverse set of classes, the goal is to identify significant differences that exceed chosen thresholds for these noise components.

A three-step process was developed to remedy the above challenges, based on the estimation of the real space noise in a given map. First, a low-pass filter is used to reduce high-frequency information in the map (low-pass filter threshold, Table 1) Clearly, this is inadvisable if high resolution is the goal for the experiment, but for heterogeneity analysis, resolution is secondary to differentiating between broader conformational and compositional differences. Second, it is important that the soft spherical mask typically applied during classification is removed, and the standard deviation of the unmasked map (σ_map_) is calculated using standard cryo-EM analysis programs. While the signal from the macromolecular object is included in this calculation, that contribution to the standard deviation is negligible if the box size is sufficiently large, so that voxels containing true signal represents 1-2% of the total map volume. Effectively, σ_map_ provides a crude estimate of the intrinsic map noise. There are several other ways to calculate a noise threshold, most recently the program developed by Beckers et al. (Beckers, Jakobi and Sachse, 2019) which uses a false discovery rate (FDR) to determine the threshold used for visualization and analysis, or one can use the noise sampled from the periphery of the map. The values of 3σ_map_ are highly correlated to the contour levels based on FDR as shown in Figure S3 but the 3σ_map_ threshold generally exceeds the FDR threshold, and is thus more conservative. Due to the prevalence of unresolved features in the ribosome data, we have used 3σ_map_ as a convenient threshold to eliminate noise. Third, each map is then binarized using a 3σ_map_ threshold such that intensities greater than 3σ_map_ were set to 1, and intensities less than 3σ_map_ were set to 0 (binarization threshold, Table 1). Other thresholds could be devised and implemented, as long as they are applied consistently across classes. These thresholded, binarized maps are used for the remainder of the analysis. Using these maps is advantageous because the “noise” from flexible regions is removed from the map, and there is a clear boundary of which parts of a structure are analyzed. Further, binarization facilitates coarse-grained analysis and eliminates the need for scaling.

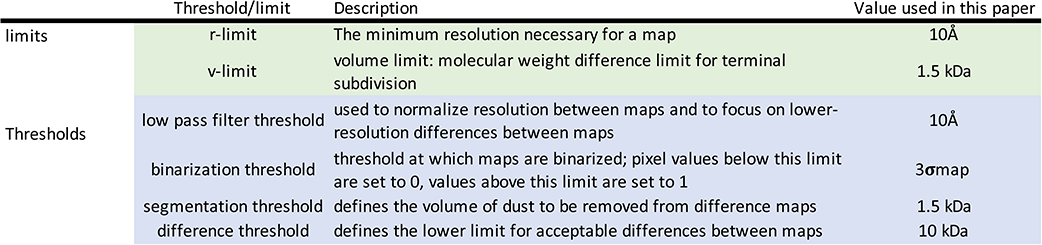

To define the endpoint to classification, the above filtering and binarization steps are applied after a test n=2 subclassification of a given class X into class X1 and X2. If either class X1 or class X2 do not have a resolvable map, as defined by the resolution limit (r-limit, Table 1), then the classification process is terminated. If the differences between class X1 and class X2 are less than the volume limit (v-limit, Table 1), the subclassification is terminated. However, if the differences between class X1 and X2 are greater than the v-limit, then the subclassification is retained, and classes X1 and X2 are in turn further subdivided into 2 classes. This process then repeats on all classes until either the r-limit or the v-limit are reached. This set of limits provides a consistent and quantitative basis for iterative subclassification.

### Segmented difference analysis between map identifies the exact number of structural states

Having discovered the structural variants in the data, we then asked how the different maps compare to one another and where/what are the major differences. To address this question, we developed a strategy to quantitatively assess similarities between the classes. While the classification approach in the discovery phase is designed to terminate once the structural features were no longer distinguishable using the r-limit or v-limit, this procedure does not guarantee that individual structures within the collective set of reconstructions are all distinct from one another. More specifically, a situation can arise where two similar classes emerge (from hiding) in different branches of the subclassification tree.

In the first step, difference maps are calculated between all of the binarized, thresholded maps. Such difference maps are useful to identify regions of density that are distinct between classes, and in our case, provide both qualitative and quantitative insight into structural relationships between distinct assembly intermediates. Two specific examples for distinct “D-classes” are shown in Figure 4A-B. The first two columns display two distinct maps arising from some point during classification. The raw difference maps are shown in the third column (map1-map2, red; map2-map1, blue). The approximate molecular weight of these differences is also indicated. These difference maps are then segmented to remove “dust” that may arise from minor conformational or compositional variations between maps. This dust cannot be interpreted in biological terms at the target resolution but may add up to a significant molecular weight (Table 1 segmentation threshold, Figure 4). Such difference maps can be computed for all pairwise combinations of reconstructions arising from the classification procedure.

**Figure 4.**
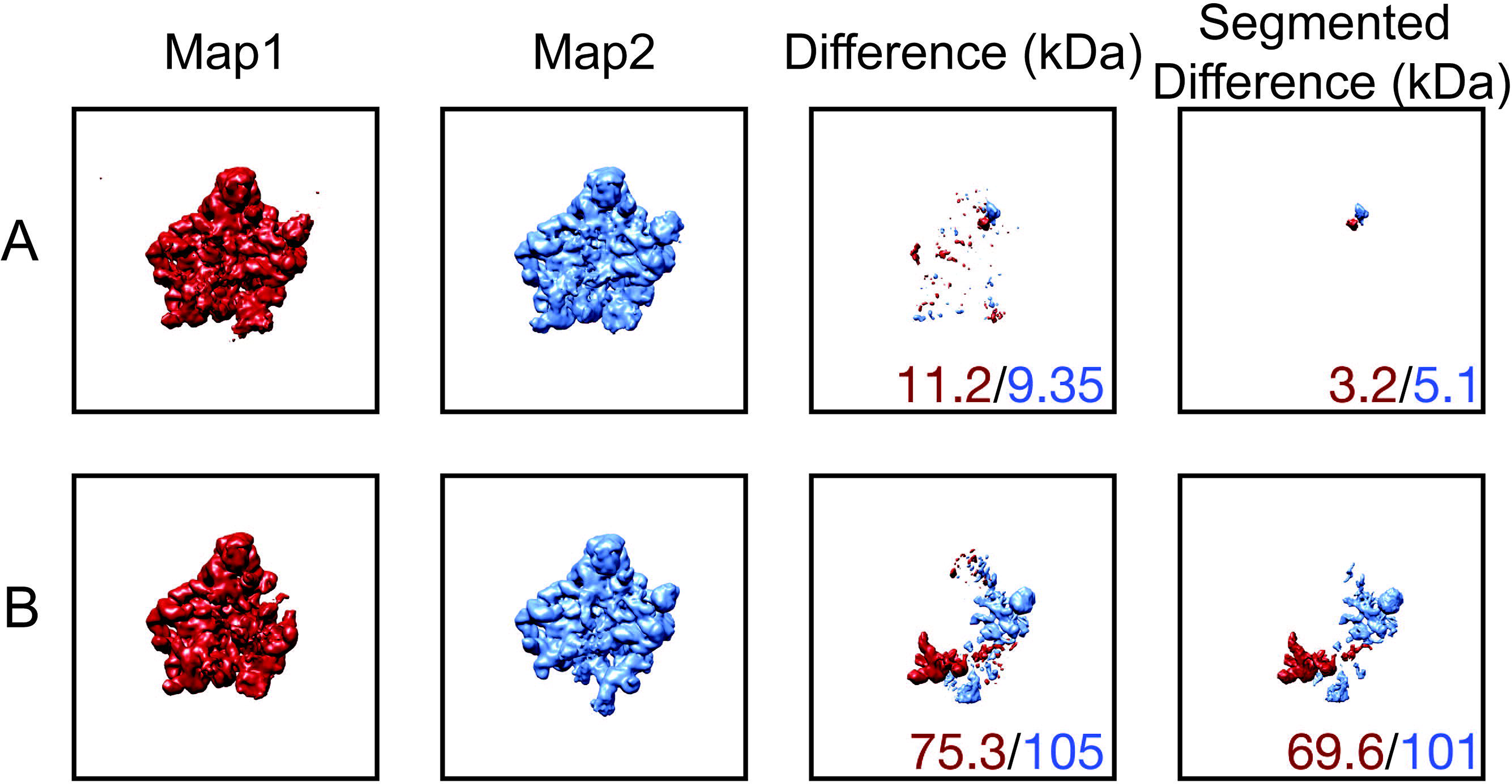
Segmented difference analysis helps to define molecular weight differences between map pairs. (A) Example where two maps would have been considered different before segmentation, but are not different after segmentation. (B) Example where two maps are different both before and after segmentation. Numbers indicate positive (Map1-Map2) and negative (Map2-Map1) molecular weight differences.

The pairwise difference maps are useful for both qualitative and quantitative downstream analyses. To parse through structural differences, define an accurate final number of *unique* structural variants in the data, and combine particles contributing to similar maps, we employed a simple hierarchical clustering approach based on the positive/negative molecular weight differences between structures. Based on the clustering, it is possible to pare down the maps and combine particles from similar reconstructions, even if they arise from different starting points in the classification (Figure 5A). At this stage, two classes can be combined if the molecular weight differences between the two classes are less than a given threshold. Since the branchpoints of the dendrogram provide a measure of *molecular weight* differences between maps, they can serve as a guide for analyzing the similarity between classes overall based on the nodes of the dendrogram (Figure 5B). In the example in Figure 5B, the dendrogram reveals that the leftmost structure is distinct from the other two and needs to be treated independently, whereas the latter two can be combined into a single class. Thus, although there are 42 distinct structures in Figure 5A, after hierarchical clustering analysis and the subsequent merging of similar maps, there are 41 distinct structures that will go forward in the analysis pathway. Collectively, these procedures enable us to identify the exact number of structural states within the data, given the limitations associated with identifying novel classes in the discovery phase and according to the established criteria in the analysis phase, defined above.

**Figure 5.**
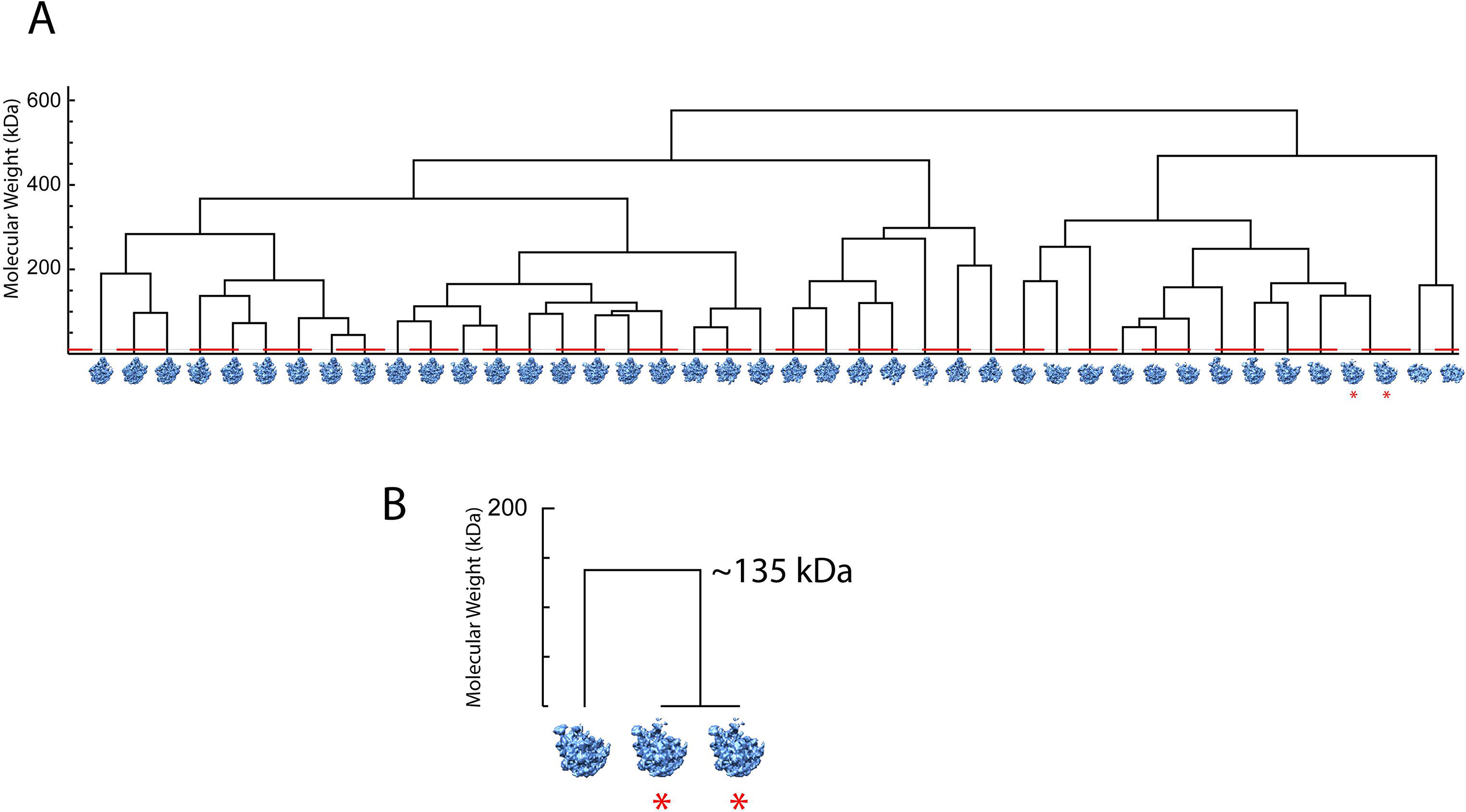
Hierarchical clustering is used to combine similar maps under a given threshold. (A) Hierarchical clustering analysis of the maps that result after the terminal subclassification (total n=42). The red dashed line indicates the 10kDa MWCO used to combine similar maps at this step, and the red stars indicate maps that are combined after this analysis. After combining similar maps, the final number of classes is thus 41. (B) Close-up example of two combined maps in (A). The leftmost structure is distinct from the other two by ~135 kDa and needs to be treated independently, whereas the two rightmost structures can be combined into a single class.

### Defining relationships between distinct structures

An important step in analyzing differences between classes discovered within the above procedures for heterogenous cryo-EM data analysis is to define *where* differences between two maps are located. If a model (e.g. an atomic model or a cryo-EM structure) exists as a reference, and if the reconstructed maps differ primarily by compositional variation, then it is straightforward to use the model for interpreting the collective set of maps (Davis *et al.*, 2016) in an “occupancy analysis.” In the case of bacterial ribosome assembly, we have a well-defined reference model (Figure 6A). This reference structure is broken into its individual r-RNA and r-protein parts, yielding theoretical cryo-EM densities for each component (Figure 6B). Such individual densities can then be directly compared to densities arising from experimental cryo-EM classification. It is important that the reference densities are generated at (approximately) the same resolution as the experimental densities arising from hierarchical clustering and difference analyses. The theoretical maps are then binarized, which enables comparing the theoretical maps to the binarized experimental maps arising from subclassification. Each binarized class (Figure 6C) is then compared to each theoretical feature map by counting overlapping voxels and normalizing to the theoretical volume, to define the fractional occupancy of the selected feature in the map that can be completely present (Figure 6D), partially present due to partial flexibility or a misdocked figure (Figure 6E), or completely missing (Figure 6F). The complete set of fractional occupancies are given as an *n* by *m* matrix of values between 0 and 1, where *n* describes the set of classes and *m* defines the number of features.

**Figure 6.**
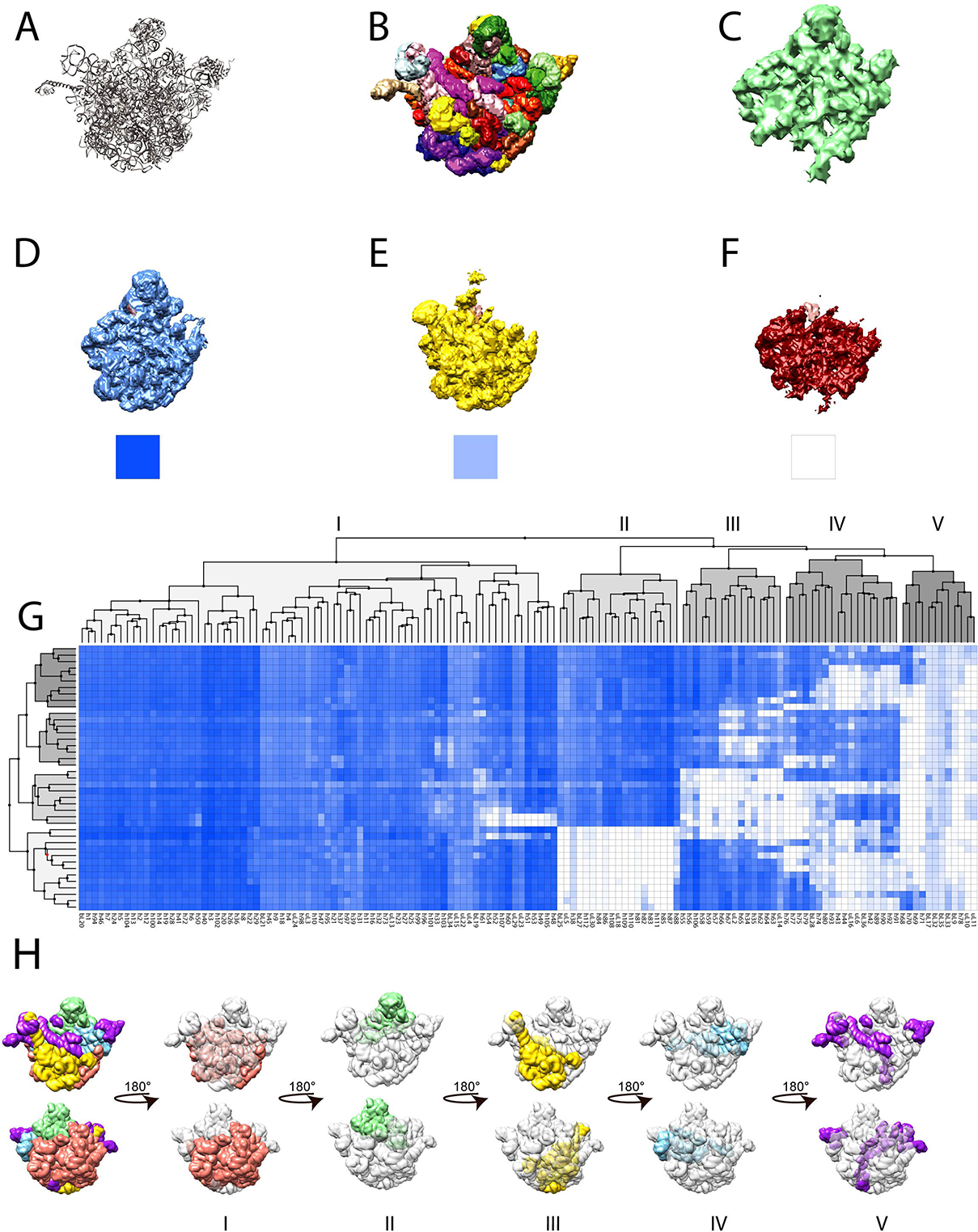
Results of occupancy analysis on the full dataset mapped onto the ribosomal scaffold (A) Reference crystal structure 4ybb. (B) Binarized maps of the individual proteins and rRNA helices created by segmenting the crystal structure into 139 individual helices and proteins, and calculating theoretical 10A maps in Chimera. (C) An example of a binarized experimental map arising from sub-classification. The pixels from the binarized experimental map that are located in the theoretical binarized map are counted and normalized to an occupancy value of 0-1. (D) Example of an E class (blue) where the occupancy of an rRNA helix (h82, salmon) is fully occupied, with the corresponding occupancy block underneath. (E) Partial occupancy example of h82 (salmon) with a C class (yellow). (F) Example where rRNA (h82, salmon) is missing in the experimental data (B class, red). G. Occupancy analysis plot, where the individual proteins and helices are shown on the x-axis, the experimental maps are on the y-axis, and the normalized occupancy values are shown from white (0) to dark blue (1). Hierarchical clustering of both structure elements and experimental maps was performed on the occupancy matrix using a squared Euclidean distance metric and Ward’s linkage. (H) Occupancy analysis blocks mapped back to the reference structure 4YBB, and the numbering system is the same as in (G).

The resulting fractional occupancies can be visualized as a heat map and subjected to hierarchical clustering to organize the classes and features (Figure 6G). Clustering along the feature (x-axis) groups elements (in this case, r-proteins and rRNAs), and clustering along the map (y-axis) groups the maps according to their occupancy. As expected, the B, C, D, and E, maps cluster well together. The occupancy matrix facilitates the visualization of large blocks of structural features that co-vary across the particle classes, providing cooperative folding blocks (Figure 6H) (Davis *et al.*, 2016). This procedure enables a quantitative comparison of distinct sets of maps that differ by compositional variants. We note that this procedure is not currently compatible with conformational variability or density that is not represented in the reference. However, if there are multiple reference models that differ by discrete conformational changes, the current protocol can be extended to competitively compare occupancies against different reference models.

### Ordering structures in a ribosome assembly pathway

In the final step that is relevant to defining an assembly process, we developed a module that uses molecular weight differences to place ribosome assembly intermediates into a pathway. In this analysis, a “folding” matrix is calculated from the molecular weight difference that would need to be added to a given map to create a second map, and the “unfolding” matrix is calculated from the molecular weight that would need to be subtracted from one map to create a second. Each element of the folding/unfolding matrix can be considered as the driving force/barrier for a structural transition between two classes. By postulating that folding proceeds by incremental assembly, with minimal unfolding, a parsimonious transition graph can be constructed with allowed passages between classes based on simple criteria – there is a molecular weight cutoff unfolding transitions, and there is a limit set to the number of transitions emanating from each class. Large unfolding events are unlikely, given the large number of states that are close in molecular weight, but small unfolding events must be permitted to allow for structural rearrangements required to transition between classes. Finally, it is likely that structural transitions proceed from a finite manifold of close intermediates. The folding and unfolding matrices can be used to construct a directed graph of allowed transitions using these criteria, as shown in Figure 7.

**Figure 7.**
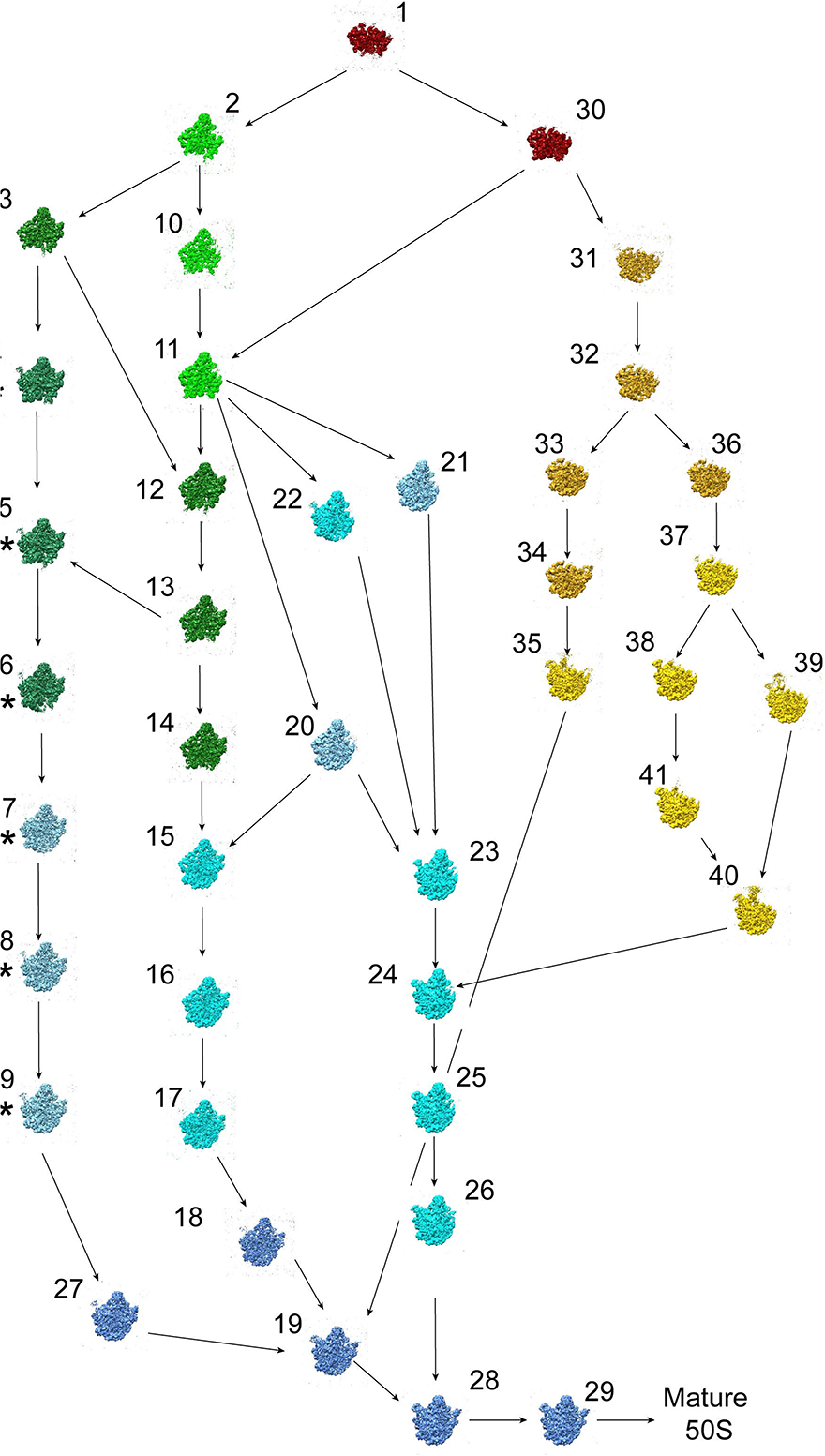
Revised ribosome assembly map from bL17-lim. (Assembly pathway drawn by analyzing the folding and unfolding molecular weight matrices and revised by hand).

### Analysis of bL17-lim data using the quantitative heterogeneity mining protocol

Our quantitative mining protocol was developed using data collected from newly purified assembly intermediates from the previously characterized bL17-limitation strain (Davis *et al.*, 2016). We collected new cryo-EM data (Supplementary Table 1 and subjected it to our workflow. In the discovery phase, we employed an updated high-resolution limit for refinement (*res_high_class=*20Å) and an initial n5>n3>n2 hierarchical classification scheme, followed by additional rounds of binary subdivision. All maps were binarized according to the 3σ_map_ threshold determined individually for each map. To determine if subclassification was complete, we selected a v-limit of 1.5 kDa. The rationale for this choice is that 1.5 kDa represents the size of the smallest RNA helix present in the bacterial ribosome and therefore corresponds to the smallest feature that we would like to capture in the data. For our purposes, smaller features can be assumed to be either biological and/or experimental noise. After iterative subclassification, the total number of classes is 42. The similarity between all of the maps was then analyzed by the hierarchical clustering analysis as described above, and at this stage, pairs of classes were combined with a 10 kDa difference threshold (Figure 5A, dotted red line). This cutoff was chosen because it is close to the average molecular weight of all proteins and rRNA features, and we wished to reduce the complexity of our data. We found one pair of structures that were similar to one another according to our established criteria for biological significance, and the particles belonging to these classes were accordingly combined (Figure 5A). Thus, using this protocol, a total of forty-one ribosome assembly intermediates were identified using quantitative metrics and similarity analysis, with minimal heuristic intervention.

The classes were then subjected to an occupancy analysis to view the sets of cooperative folding blocks across the different classes. For the bL17-lim dataset here, the maps are compared to the reference crystal structure (PDB 4ybb) (Figure 6A). The reference 4ybb structure is filtered to 10Å and segmented into volumes corresponding to individual r-proteins and rRNA helices, resulting in 139 theoretical map segments (Figure 6B). The fractional analysis revealed five major structural blocks (Figure 6G,H) The largest, block I (red) is composed of structural elements that are largely present in all of the classes. These elements are found on the back of the ribosome and represent the structural core that can form without bL17. Block II (green) represents the central protuberance, which is fully formed in the D and E classes but is either missing or misdocked in the B and C classes. Block III (yellow) maps to the base of the ribosome and the L1 stalk. These features are mostly present in the C and E classes but are missing in the B and D classes. These two blocks represent parallel pathways in assembly (Davis *et al.*, 2016), as it is unlikely that the base of the ribosome would be unfolded or disordered in order to form the central protuberance, and vice versa. Block IV (blue) represents density that is specific to the base of the L7/12 stalk and is mostly present in the D and some of the E classes. Finally, block V (purple) represents density that is mostly missing in all maps, and is composed of h68, bL9, the L7/12 stalk, and the top of the L1 stalk. These represent features that are among the last of the ribosome to fold (like h68) or are flexible elements (the stalks). bL9 is a special case, as the conformation in the crystal structure is an artifact due to crystallization; in cryo-EM structures, bL9 wraps around to the interface between the 30S and 50S subunits and is often flexible. These central blocks are very similar to the ones that we discovered previously (Davis *et al.*, 2016), but this updated occupancy matrix will allow us to compare the blocks that arise from other depletion or deletion strains in order to explore the cooperative block-like behavior or ribosome assembly in future work.

The ordering module was used to calculate an initial pathway in the absence of bL17-lim, which was modified by hand, as elements like the misdocked central protuberance and non-native structural elements can have large effects on molecular weight differences but may arise earlier in the order of assembly. We found the same initial super classes as previously reported (B, C, D and E classes). While the classes we found were similar to the initial bL17 data (Supplemental Figure 4), the new classes enabled refinement of our bL17-lim ribosome assembly pathway. First, we found a YjgA-dependent pathway through the assembly process (Figure 7, classes denoted by *). YjgA was only bound if the central protuberance and the L1 stalks were present. We also discovered three potential parallel processes in the C class where the earliest event could either be the completion of the L1 stalk, the partial docking of the central protuberance, or the formation of the base of the L7/12 stalk. We did not previously observe the formation of the base of the L7/12 structure in the assembly pathway for any class. We also found an immature B class (Figure 7, structure 1) and an immature D class where the base was missing, but the L7/12 stalk was absent or present (Figure 7, structures 2 and 3), which were not present in the original set of 13 structures (Davis *et al.*, 2016). In particular, the immature B class represents the least mature pre-50S intermediate identified to date. We also identified several structures that seem to be transition points between the two classes (Figure 7, structures 6 and 7), and we observe formation of density at the base of the structures, which is lacking in other D classes and is present in other E classes. These new discoveries inform a better understanding of ribosome assembly in the context of bL17 limitation, and the data analysis process will allow us to quantitatively assess cryo-EM data from other limitation strains and ribosome assembly defects.

## Conclusions

Heterogeneity analysis in cryo-EM provides exciting opportunities to discover new biology, but current workflows suffer from numerous challenges. The work here addresses three challenges that researchers face in the analysis of cryo-EM data, as exemplified using a case study of ribosome assembly intermediates: establishing a divisive approach to classification with well-defined endpoints to discover novel stats, a comprehensive difference analysis between distinct structures, and the application of well-defined criteria (thresholds) for limiting classification. The application of specific thresholds and limits (Table 1) has been critical to the success of analyzing ribosome assembly intermediate data. The implementation of this workflow has allowed us to identify an additional 28 ribosome assembly intermediates (counting the 41 assembly intermediates after merging similar classes), which include an independent pathway for the assembly factor YjgA and the earliest intermediate discovered to date in the ribosome assembly process. The discovery and analysis modules of this workflow provide a powerful analysis for quantitatively interrogating heterogeneous cryo-EM data for complex biological processes.

## Acknowledgements

Molecular graphics and analyses were performed with the USCF Chimera package (supported by NIH P41 GM103311). This work was supported by grants from the NIH DP5-OD021396 and U54 AI150472 (to D.L.) R35-GM136412 (to JRW).

## Author Contributions

Jessica N. Rabuck-Gibbons: Conceptualization, Investigation, Methodology, Software, Formal Analysis, Data Curation, Writing – Original Draft, Writing – Review & Editing, Visualization.

Dmitry Lyumkis: Conceptualization, Investigation, Writing – Review & Editing, Resources.

James R. Williamson: Conceptualization, Methodology, Software, Data Curation, Writing – Review & Editing, Visualization, Supervision, Project Administration, Funding Acquisition.

## Declaration of Interests

The authors declare no competing interests.

## Materials and Methods

### Cell Growth and Isolation of Ribosomal Particles

Cells were grown and ribosomal particles were isolated as in (Davis *et al.*, 2016). Briefly, strain JD321 was grown in M9 media (48mM Na2HPO4, 22mM KH2PO4, 8.5mM NaCl, 10mM MgCl2, 10mM MgSO4, 5.6mM glucose, 50mM Na3*EDTA, 25mM CaCl2, 50mM FeCl3, 0.5mM ZnSO4, 0.5mM CuSO4, 0.5mM MnSO4, 0.5mM CoCl2, 0.04mM d-biotin, 0.02mM folic acid, 0.08mM vitamin B1, 0.11mM calcium pantothenate, 0.4nM vitamin B12, 0.2mM nicotinamide, 0.07mM riboflavin, and 7.6mM (14NH4)2SO4]) with tetracycline (10 mg/mL), chloramphenicol (35 mg/mL),and limiting conditions HSL (0.1 nM) and harvested at OD=0.5. Cells were lysed in Buffer A (20mM Tris-HCl, 100mM NH4Cl, 10mM MgCl2, 0.5mM EDTA, 6mM b-mercaptoethanol; pH 7.5) by a mini bead beater, and the clarified lysate was fractionated on a 10-40% w/v sucrose gradient (50mM Tris-HCl, 100mM NH4Cl, 10mM MgCl2, 0.5mM EDTA, 6mM b-mercaptoethanol; pH 7.5).

### Electron Microscopy Data Collection

Fractions containing the ribosomal intermediates were spin-concentrated with a 100 kDa MW filter (Amicon) and buffer exchanged into Buffer A. 3 μl of this sample was added to a plasma cleaned (Gatan, Solarus) 1.2mm hole, 1.3mm spacing holey gold grids (Russo and Passmore, 2014). Grids were manually frozen in liquid ethane, and single particle data was collected using Leginon on a Titan Krios microscope (FEI) with a K2 summit direct detector (Gatan) in super-resolution mode (pixel size of 0.66Å at 22,500 magnification). A dose rate of ~5.8e^-^/pix/sec was collected across 50 frames with a fluence of 33-35e^-^/Å^2^ at a tilt of -20° to compensate for preferred orientation (Tan *et al.*, 2017b).

### FrealignX Classifications

After conversion from Relion to FrealignX parameters, global refinements were performed in FrealignX, and all occupancies were randomized across the parameter files. A final value of 20Å was selected for *res_high_class*, and after every 10 cycles of classification/refinement, all classes were aligned to a C class scaffold using custom scripts for a 3D alignment with Chimera (Pettersen *et al.*, 2004) while running FrealignX. For each classification step, 50 refinement/classification cycles were performed. After initial classification, each class was selected in a parameter file for subsequent rounds of classification using the merge_classes.exe in cisTEM (Grant, Rohou and Grigorieff, 2018) and custom scripts. The occupancies were randomized across the parameter files, and the same cycle of 50 cycles of refinement/classification interspersed with 3D alignment with Chimera every 10 cycles. FSC curves and Euler plots were generated by FrealignX and cisTEM (Grant, Rohou and Grigorieff, 2018), and 3DFSC plots were calculated by the 3DFSC server (Tan *et al.*, 2017a). The SCF was calculated according to the process in (Baldwin and Lyumkis, 2021; Baldwin and Lyumkis, 2020). The 3DFSCs and all maps shown were visualized in Chimera (Pettersen *et al.*, 2004), and the details for each map are indicated in Table S1.

### Calculation of σ values

For analysis, each map was first filtered to 10Å. To calculate σ which was used as a measure of noise, each map was unmasked by expanding the *outer_radius* in FrealignX so that the spherical particle mask would be larger than the box size. The Fourier folding of signal along the edges of the box was negligible. Relion 2.1 was used to calculate the σ value using the *relion_image_handler* command. Relion 2.1 was then used to create binarized maps using the *relion_image_handler* command, and the binarization threshold was set to 3σ.

### Hierarchical clustering analysis

Thresholded, binarized maps were given as input to a custom Mathematica script (Wolfram Research, 2020). The Mathematica script calculated the segmented difference maps between all maps and calculated the molecular weights of the differences maps (in kilodaltons) using Equation 1(Ludtke, 2016):

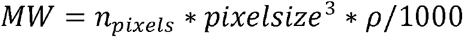

Density ρ is 0.81 daltons/Å^3^. The MW difference matrix was clustered using the Euclidean distance metric and Ward’s linkage and displayed in a dendrogram. Similar maps were averaged together after hierarchical clustering analysis using EMAN2 ((Ludtke, 2016).

### Occupancy Analysis

The thresholded and binarized maps were given as input, and the reference map from the *E. coli* 50S subunit crystal structure (PDB ID 4YBB) was segmented into 139 elements comprised of individual ribosomal proteins and rRNA helices according to the 23S secondary structure. Theoretical densities for each r-protein and rRNA helix were calculated for each element at 10Å using the pdb2mrc command from EMAN. Prior to binarization, voxels that had overlapping theoretical density from two structural elements, were assigned to the smaller of the two theoretical volumes so that each pair of volumes is nonoverlapping. Each voxel density was binarized to either 0 or 1 using a threshold of 0.016, which is the threshold that gave the approximately correct molecular weight for individual r-proteins and rRNAs helices. The relative volumes in the binarized experimental and reference maps were calculated, which gave a fractional occupancy between 0 and 1 for each element. The occupancy values were clustered across the rows (classes) and columns (rRNA/protein elements) using an unsupervised hierarchical clustering using the Euclidean distance metric and Ward’s linkage method, as implemented in Mathematica.

### Parsimonious folding/unfolding matrices

A pathway diagram was constructed by using the *n* × *n* molecular weight difference matrices, **M**_**f**_ and **M**_**u**_, from a set of *n* structures. Each difference map (M_i_-M_j_) has negative elements corresponding to folding that occurs in the transition from class i to class j, and positive elements that correspond to unfolding that occurs in the transition from class i to class j. The volume changes for folding and unfolding form the elements of **M**_**f**_ or **M**_**u**_, noting that **M**_**u**_ = **M**_**f**_^**T**^. The matrices **M**_**f**_ and **M**_**u**_ are used to construct a directed graph **G**, comprised of the set of vertices *v_i_*, and a set of directed edges, *e_ij_*, representing the allowed transitions between classes, The set of edges is initialized as the set of *e*_*ij*_ where M_u,i,j_ > M_f,i,j_, such that only net folding transitions are allowed. The set of edges is pruned using two global parameters: θ_unf_ as a maximum threshold for unfolding, and n_branch_, as a limit on the number of transitions emanating from a single class. The unfolding threshold limits unreasonable structural rearrangements, while the branching threshold limits transitions to a small set of the closest transitions. Edges are eliminated if the unfolding exceeds the threshold such that M_u,i,j_ > θ_unf_, unless elimination of the edge results in a disconnected graph **G**. Next, for each vertex *v_i_*, the set of remaining edges *e*_*ik*_ emanating from *v*_*i*_, are sorted into the order based on the M_f,i,k_, retaining at most the n_branch_ edges, again, unless deleteing the edge would result in a disconnected graph **G**. The resulting transition graph **G** should have one or more *source* vertices (classes) that are the earliest classes in the assembly pathway, and one or more *sink* vertices that are the most mature classes in the pathway. Tuning of the parameters θ_unf_ and n_branch_, adjusts the connectivity and degree of branching of the resulting graph. The graph vertices are annotated with thumbnails of the map, followed by manual layout of the graph into a sensible order in Adobe Illustrator. The values of θ_unf_ and n_branch_ used to generate the graph in Figure 7 were 390 kDa and 3, respectively.

### Data Deposition and Software Availability

Mathematica scripts and example parameter files, where needed, will be available upon request. All maps are deposited at EMPIAR as noted in the Key Resources Table.

## Supplemental Information Titles and Legends

Supplemental Figure 1. Results of the five tested classification schemes grouped by class. The structures are colored by classification scheme. Unique classes are shown by an asterisk (*), and classes that are similar are underscored by red brackets. Clustering of (A) the B classes, (B) the C classes, (C) the D classes, and (D) the E classes resulting from the tested classification schemes. Any 70S or “junk” classes that result from the subclassifications are omitted for clarity.

Supplemental Figure 2. Example of ambiguous density and features for B class particles. From left to right: (B class filtered to 5Å and shown at 3σ_map_, 2σ_map_, and 1.5σ_map_. In particular, at the 2σ_map_ threshold, noise above background is visible proximal to the main particle that is likely due to disordered rRNA (black arrows).

Supplemental Figure 3. Comparison of the the 3σ_map_ threshold that is used in our current analysis versus the confidence map FDR threshold (Beckers, 2019). The black line represents y=x, and the red and black dots represesent thresholds at 1% and 0.01% FDR, respectively. The measures are highly correlated, and the 3σ_map_ threshold is generally more conservative than either FDR threshold.

Supplemental Figure 4. Hierarchical clustering analysis of the original bL17-lim data (orange) together with the structures solved by the new data processing workflow (blue). The red dotted line indicates the 10.0 kDa cutoff applied to determine similarity between classes. The original classes typically have counterparts within the new data (red underlined structures), but the new workflow is able to identify many more structural intermediates.

## Notes

### Competing Interest Statement

The authors have declared no competing interest.

