## Supplementary figures and images for "Quantitative Mining of Compositional Heterogeneity in Cryo-EM Datasets of Ribosome Assembly Intermediates"

### Supplemental Figure 1

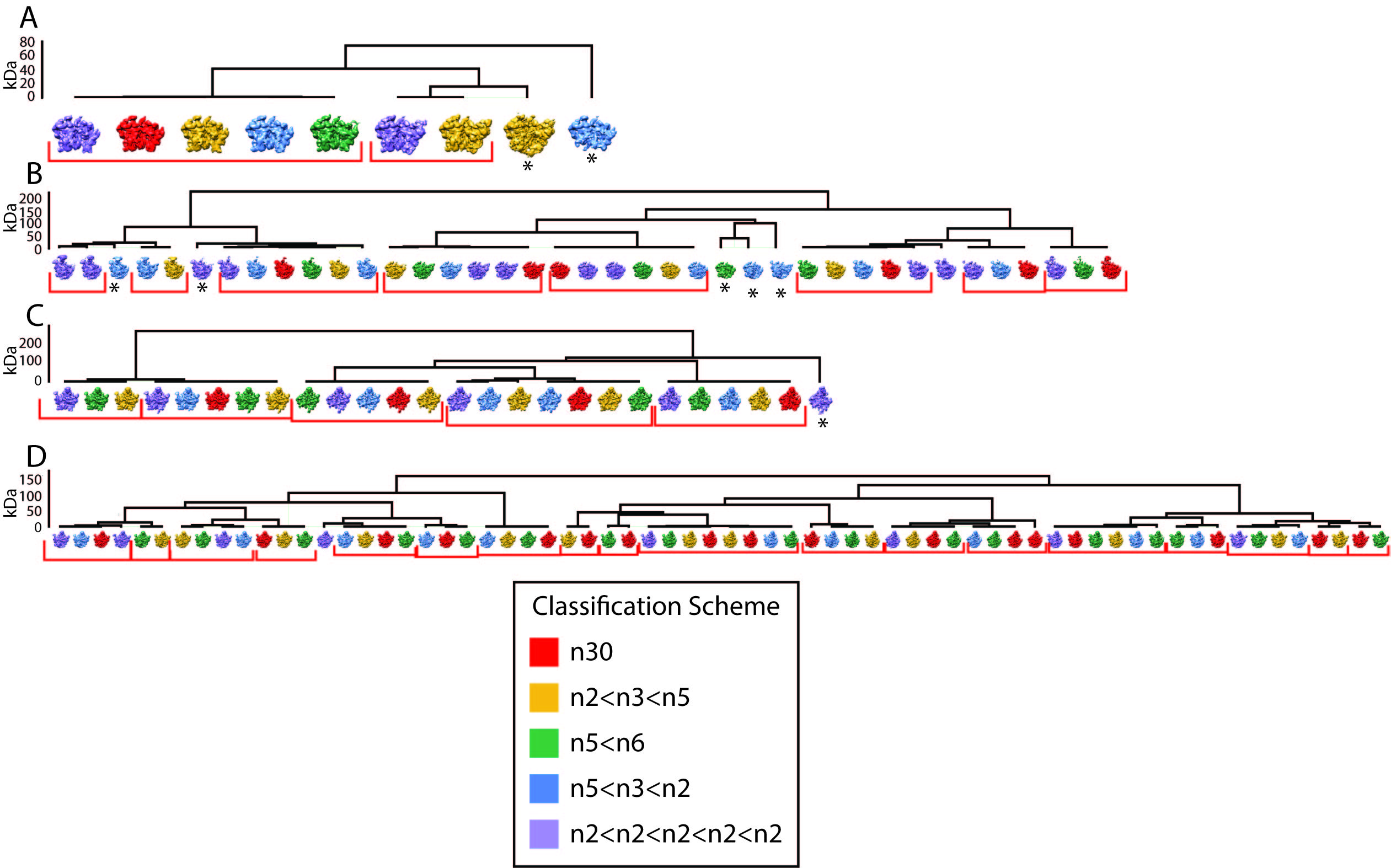

### Supplemental Figure 2

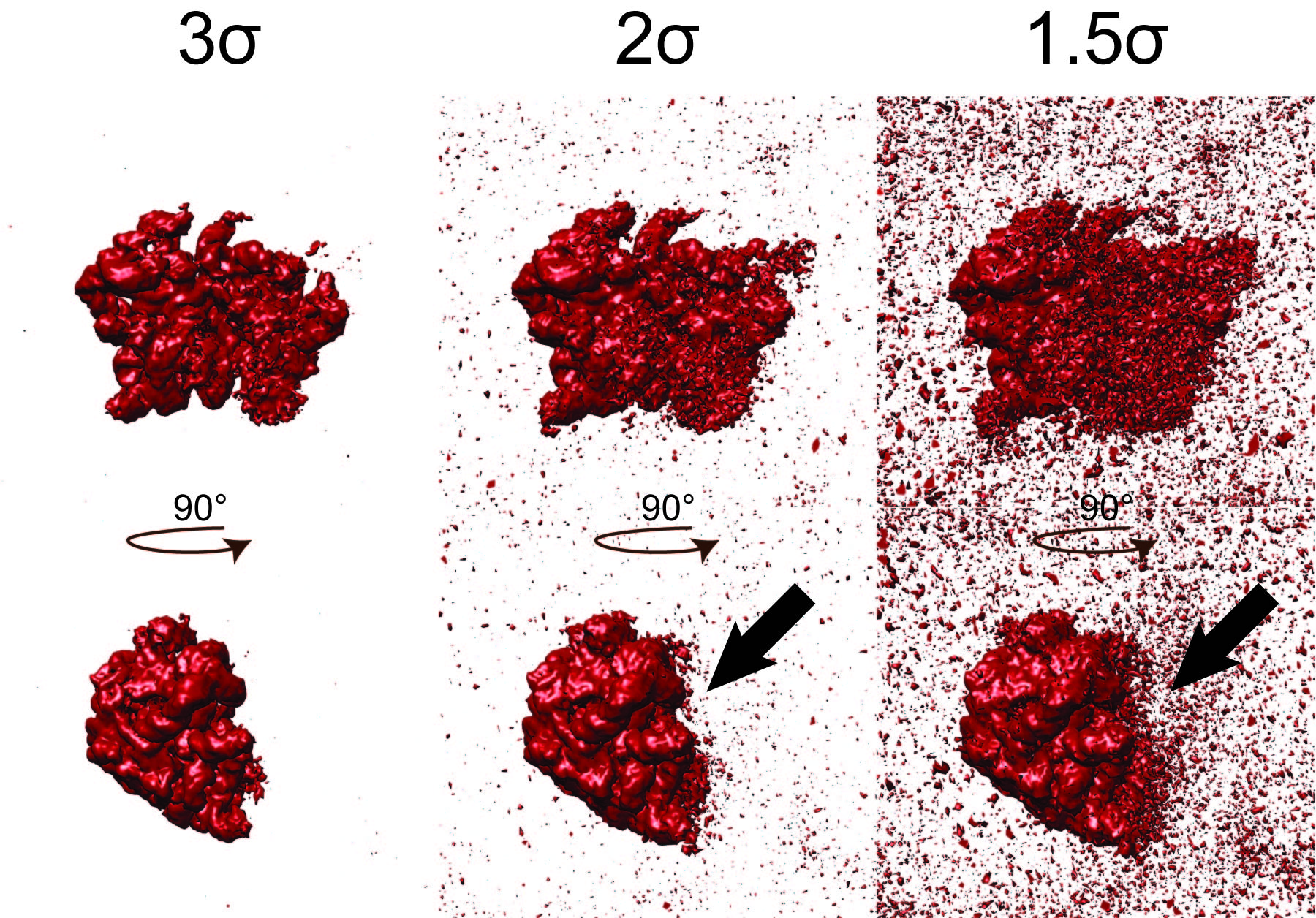

### Supplemental Figure 3

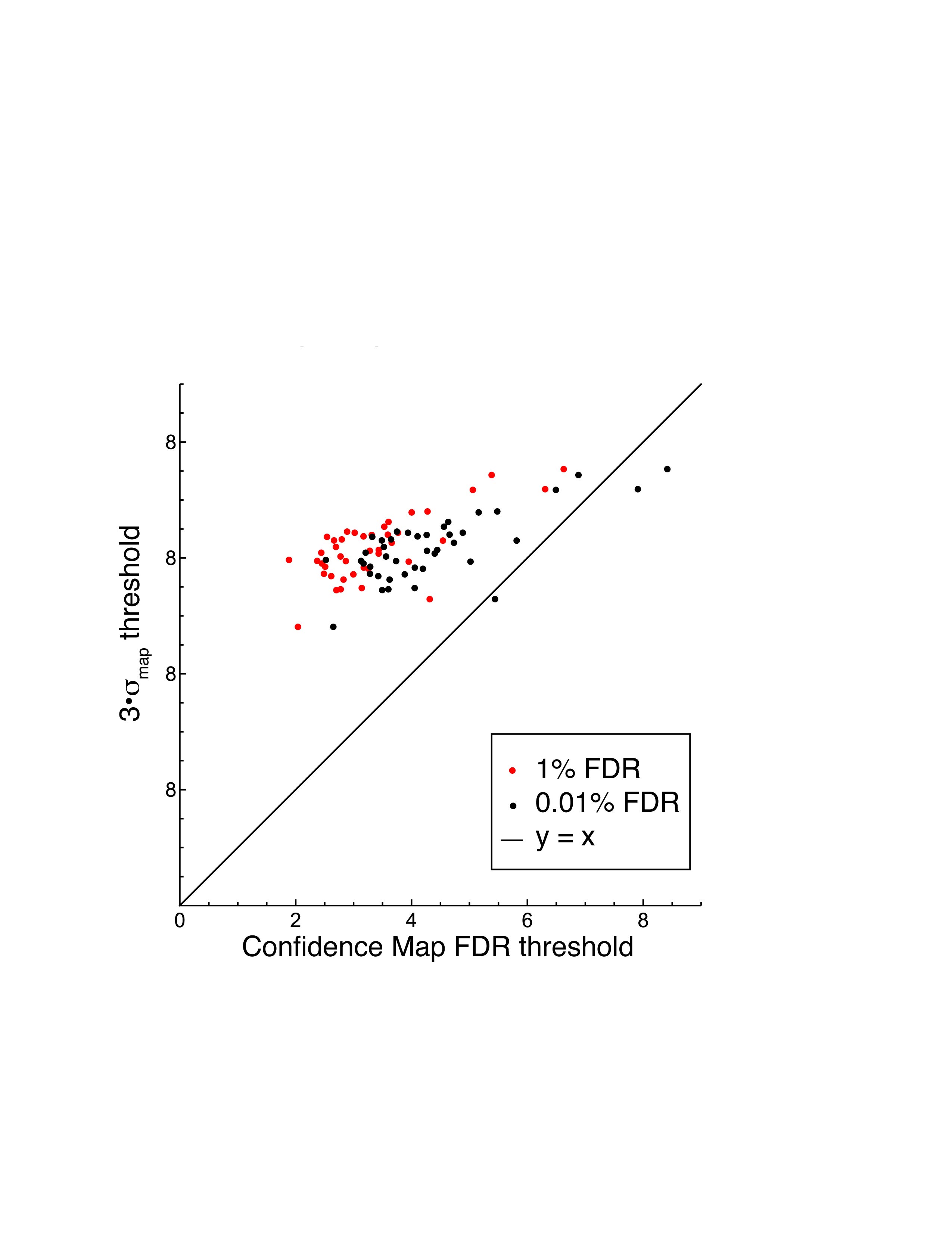

### Supplemental Figure 4

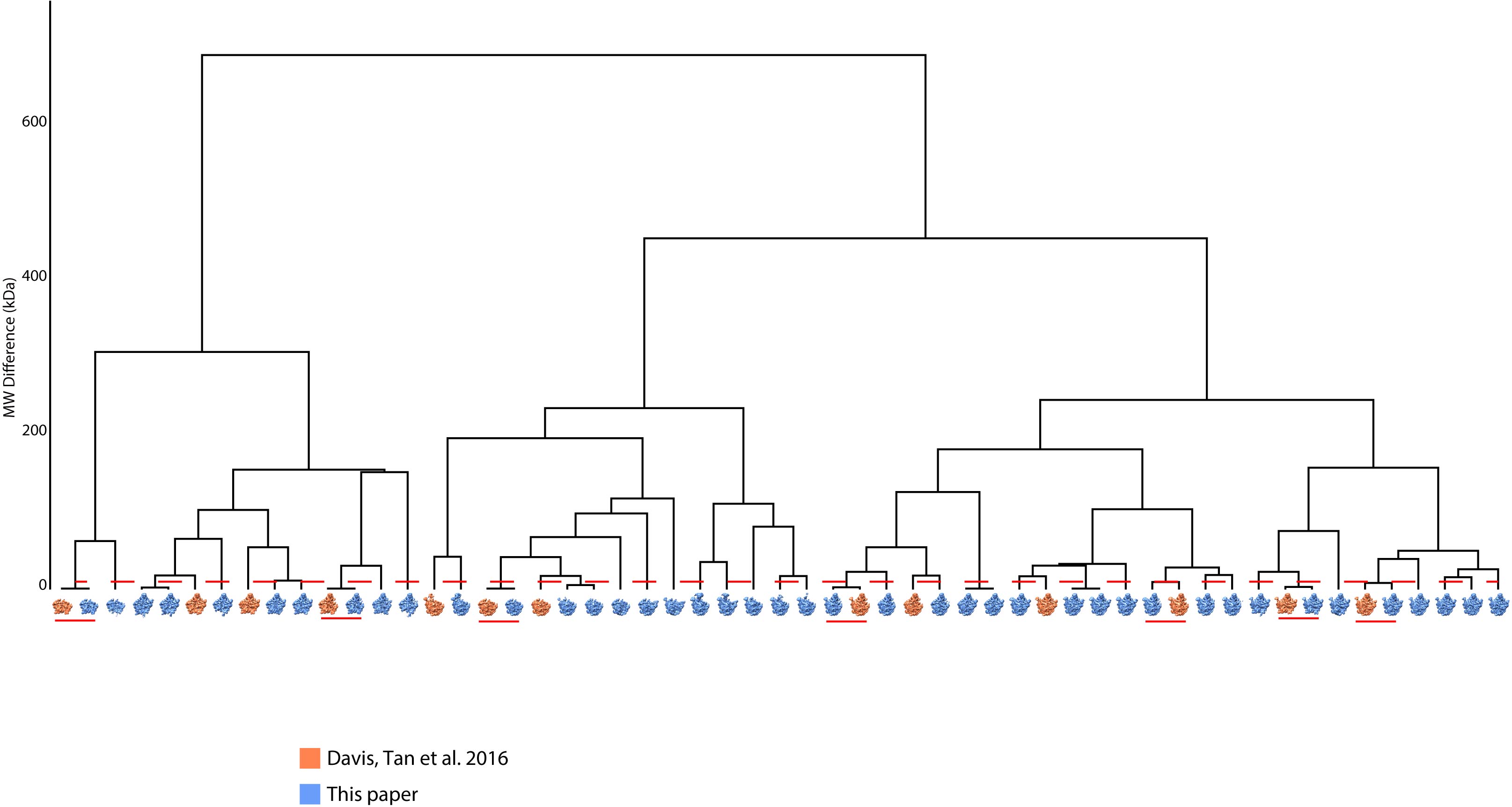
